# Characterizing the 3D structure and dynamics of chromosomes and proteins in a common contact matrix framework

**DOI:** 10.1101/264515

**Authors:** Richard J. Lindsay, Bill Pham, Tongye Shen, Rachel Patton McCord

## Abstract

Any conformational ensemble of biopolymers, whether they are proteins or chromosomes, can be described using contact matrices derived from experimental or computational studies. One powerful approach to extract meaningful information from these contact matrices is to perform principal component analysis (PCA) on the covariance matrix of the contact data. Indeed, PCA on Hi-C chromosome contact matrices has revealed the spatial segregation of active and inactive chromatin. Separately, PCA on contact matrices from snapshots of protein conformations has characterized correlated fluctuations of protein domains. However, despite the similarities of these data and analyses, there has been little synergy between the PCA approaches and the comparison of resulting biological insights obtained for protein and chromosome structures. We note that, to date, different styles of analyses were applied exclusively to each biomolecule type: explicit contact correlation analysis (E-PCA) for proteins and implicit contact correlation (I-PCA) for chromosomes. In this work, we compare the results of applying both methods to both classes of biopolymers. While I-PCA reveals only average features of 3D chromosome structure, we find that applying E-PCA to an ensemble of chromosome structures from microscopy data reveals the dominant motion (concerted fluctuation) of the chromosome. Applying E-PCA to Hi-C data across the human blood cell lineage isolates the aspects of chromosome structure that most strongly differentiate cell types. Conversely, when we apply I-PCA to simulation snapshots of two protein complexes, the major component reports the consensus features of the structure, while the previously applied E-PCA characterizes correlated deviations from the mean structure.

## Introduction

The functions of large classes of biopolymers are related to their structural stability and conformational dynamics, from small scale conformational changes of proteins responding to chemical and physical stimuli to large scale genome structure reorganization. While the 3D structure of biopolymers is often represented by the Cartesian coordinates of each point along the polymer, other structure representations, such as contact matrices, can facilitate analyses of the configuration and motion of biopolymers in many situations. The structural features of a folded polymer can be captured by recording the contacts formed between different parts of the molecule. This information can then be organized in a contact matrix in which the linear constituents (amino acid residues or genomic positions) of the polymer are labeled along the rows and columns of the matrix.

In protein studies, such contact information is typically derived from a 3D structure (obtained from either experiments and/or computer simulations), defining a “contact” when one region of the amino acid chain is within a certain distance of another region (1, 2). For chromosome structure, a similar distance threshold approach can be employed when high resolution microscopy data (showing the path of a chromosome in 3D space) is available (3). Even more commonly, however, chromosome contact matrices are measured directly using chromosome conformation capture experiments (4). This approach chemically captures contacts between chromosome regions using formaldehyde crosslinking followed by DNA digestion and proximity ligation. Then, in the genome-wide version of the technique, Hi-C, contacts are identified by high throughput sequencing of ligated DNA pairs (5). In most Hi-C experiments, the resulting chromosome contact matrices report the frequency of contacts between pairs of chromosome loci within a population of cells. Single cell Hi-C techniques are also emerging (6), resulting in contact maps at single cell resolution, which are directly analogous to the protein structure contact maps described above.

Contact matrix descriptions of polymer configurations, and the ensuing statistical analyses of the matrices, have proven to be tremendously useful for the study of protein and chromatin structures. The first chromosome contact matrices generated by Hi-C immediately revealed major principles of chromosome folding. In particular, a "plaid" pattern on the contact map was found to represent the spatial compartmentalization of active and inactive regions (also termed the A and B compartment) along the chromosome(5). This pattern can be mathematically isolated and quantified using a version of Principal Component Analysis (PCA)(7) in which each row of the contact matrix is treated as a degree of freedom (details described below) (5). As increasing numbers of Hi-C contact matrices are published in different cell types and conditions, PCA has continued to be frequently used to analyze this population average chromosome spatial compartmentalization (8, 9). Meanwhile, PCA has been extensively used to characterize protein conformational dynamics, especially directly using the Cartesian coordinates of the system (10–12). In recent years, PCA has also been used for the statistical analysis of other degrees of freedom (DOFs), such as torsion angles (13) and residue-residue contacts (14) in the protein system. Particularly, treating each individual contact as a DOF, PCA assisted researchers in identifying regions of protein with concerted dynamics of contact forming and breaking (14–20).

Even though the underpinning mathematics of protein and chromosome contact matrices is identical, because protein and chromosome structure analyses have largely been developed in separate communities of researchers thus far, there has been little comparison of the properties of these contact matrices. No connection between different approaches to similar analyses (such as PCA) has been made. A unified viewpoint and comparison of the analyses performed using contact matrices in these different biological systems may facilitate the further development of these research areas. Here, we formally describe the differences between the divergent contact correlation analyses that have been used to date exclusively on either proteins or chromosomes, and seek to determine the advantages, disadvantages, and different biological insights about protein and chromatin structure that can be determined by each method. We term the PCA method previously applied exclusively to chromatin contact matrices “implicit contact correlation analysis” (I-PCA) and the method previously applied exclusively to proteins “explicit contact correlation analysis” (E-PCA).

Both I-PCA and E-PCA begin with contact matrices of size N × N (for a polymer comprised of *N* constituents: amino acids, or genomic bins, for example). Once contact matrices are obtained, a covariance matrix of the contact variables is calculated. Then, PCA of the covariance matrix reveals major properties of the contact pattern. The essential difference between E-PCA and I-PCA is how the initial stochastic contact variables are defined. In E-PCA analysis, explicit contacts (contacts between two explicitly labeled parts of the biopolymer, *i* and *j*) are treated as independent variables. When a single conformation can be specified, as often is the case of protein data input, the value of can be simply assigned to either 1 or 0 depending on whether a contact is made. Other more elaborate improvements include using continuous contact energy (18) and coarse-graining the contacts (19). For genome Hi-C structures, in the absence of single cell data, the value of *u_ij_* is the number of cross-links measured with an elegant normalization procedure (5). E-PCA tracks the correlation between these *N*^2^ contact elements *u_ij_* (as shown in the dashed red square in Figure 1) across *T* snapshots or samples, and the size of covariance matrix is *N^2^ × N^2^* in principle, as illustrated in Figure 1. Practically, due to symmetry, only the *N* × (*N* + 1)/2 unique contacts are used. Thus, the covariance matrix of E-PCA explicitly provides correlation information between four parts of the biopolymer, *C_ijkl_* = 〈(*u_ij_* – 〈*u_ij_* 〉) ( *u_kl_* – 〈 *u_kl_* 〉) 〉 that is, when *i* and *j* form a contact, whether *k* and *l* are likely to form a contact (Figure 1). Here 〈 〉 indicates an ensemble average.

On the other hand, the stochastic contact variables of I-PCA analysis, *u_i_*, are whole rows (or columns) of the contact matrix, as shown in the red rectangle of Figure 1. That is, for polymer with *N* units, one has total *N* stochastic variables. The contact variable *u_i_* is the pattern of contacts between *i* and the rest of the polymer. Each contact matrix has *N* rows and thus contributes *N* sample points, comparing to the fact that in E-PCA, each entire contact matrix contributes to 1 sampling point. The covariance matrix of I-PCA, *C_ij_* = 〈( *u_i_* – 〈*u_i_*〉) (*u_j_* – 〈*u_j_*〉)〉, reveals the correlation between the pattern of contacts between *i* and the rest of the polymer vs. the interaction pattern of *j* with the rest of the polymer (Figure 1). In summary, for a total *T* contact matrices of size *N* × *N*, E-PCA identifies N^2^ contact variables (more precisely, *N*(*N* + *1*)/*2*) and total *T* sampling data points while I-PCA identifies total N contact variables and total *T × N* data points. The variation of the I-PCA scheme that has been previously applied to the population-averaged Hi-C contact maps, we term such a procedure "mean implicit contact correlation analysis" (MI-PCA). If the contact matrices are not already an average (as is often the case with Hi-C data), one first obtains the mean contact matrix of *T* matrices and then performs I-PCA on this mean matrix. It should be stressed that MI-PCA has *N* contact variables and only *N* data points.

In this article, we will first present the application of E-PCA to contact matrices of human chromosome 21 obtained from microscopy data (3). Interestingly, this study obtained detailed snapshots of individual chromosome structures from many different cells, data which is very similar to the protein structure snapshots previously analyzed by E-PCA. But, the only PCA analysis performed in the original analysis of this data was MI-PCA (as usually performed on Hi-C maps) on the mean spatial distance matrix to show that spatial compartments revealed by imaging matched previous reports from Hi-C data. This reiterates the observation that, while commonly used to study protein structural dynamics, PCA has never been applied to analyze correlated changes in structure across heterogeneous chromosome data. Our results suggest essential correlated fluctuations of chromosome structure across imaging snapshots. We further explore the utility of the E-PCA method in chromosome structure analysis by applying it to chromosome contact matrices from Hi-C data collected across a group of related blood cell types (21). Our results show that E-PCA can highlight dominant modes of chromosome structure changes between cell types. We demonstrate the results of applying I-PCA and MI-PCA contact analyses to protein conformations using two nuclear hormone receptor complexes as examples. Applying E-PCA to chromosomes and I-PCA to proteins allows us to find analogies and contrasts between methods and between these different biopolymer systems. And, to our knowledge, it is the first time such analyses have been performed on the corresponding systems.

## Results

### Explicit contact correlation analysis reveals correlated variations in chromosome structure between individual cells

E-PCA has previously been used to examine correlated fluctuations across many snapshots of protein structures. In the early days of 3D chromosome conformation studies, there were not enough sets of data available on different individual chromosome structures to make this analysis possible for chromosomes. Now, with increasing numbers of Hi-C datasets, single cell Hi-C, and high resolution microscopy data, we can apply E-PCA to chromosome structure data. The essential difference between E-PCA, which we now apply to chromosome structure data, and I-PCA, which has been previously applied to chromosome contact matrices, is the number of contact degrees of freedom. As shown in the cartoon illustration shown in Fig. 1, in a 4×4 matrix, I-PCA examines the correlation between four contact variables, each representing the contact pattern of one biopolymer element (ie genomic bin) with the rest of the polymer. E-PCA, on the other hand, examines the correlation between sixteen contact variables across many related contact matrices, where each variable is an explicit contact between two sites along the biopolymer. For an E-PCA analysis, one first obtains contact correlation matrix *C_ijkl_* = 〈( *u_ij_* – 〈*u_ij_*〉) (*u_kl_* – 〈*u_kl_*〉)〉, and then proceeds with principal component analysis. Since the computational complexity of obtaining *C_ijkl_* is in the order of *0(N^4^)*, and the computation time increases in proportion to the quartic power of the system size *N*, we also term E-PCA a four-point correlation analysis.

**Figure 1:**
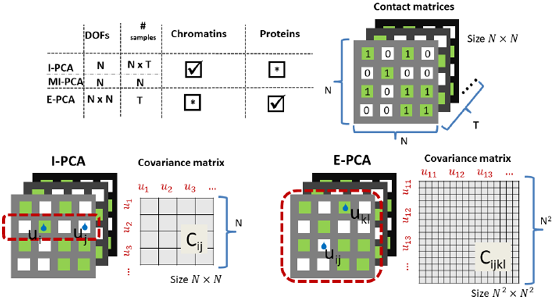
An illustration of contact correlation analysis methods. Here, identical input information, an ensemble of contact matrices, is being digested two different ways. In explicit contact correlation analysis (E-PCA), specific contacts are treated as independent variables and their covariance matrix is calculated using the whole contact matrix as one sample (dotted red line). On the other hand, implicit contact correlation analysis (I-PCA) treats rows of the contact matrix (a contact pattern between one constituent of the polymer and the rest of the polymer) as independent variables and calculates the covariance matrix using each row as one sample (dotted red line). In the table, check symbols indicate the previous pairings of analysis methods and systems while stars indicate the new pairings explored in the current work.

For the first application of E-PCA to chromosome structure, we re-examined an ensemble of structural data on human chromosome 21 (Chr21) obtained by high resolution microscopy (3). In this experiment, Topologically Associating Domains (TADs) identified by previous ensemble Hi-C experiments (22) were used as structural subunits along Chr21 and were labeled with fluorescent probes in IMR-90 fetal lung fibroblast cells. The resulting data provide the 3D spatial coordinates of each TAD along Chr21 in 120 myofibroblast cells (only 47 cells for which data exists for every TAD are being used here, i.e., *T* = 47). The original analysis of this data used MI-PCA on the average chromosome structure to define the "A" and "B" compartments (positive and negative elements of eigenvector PC1) and to demonstrate that active and inactive chromatin compartmentalization is detected with microscopic chromosome tracing just as it is found with ensemble average Hi-C data. One compartment ("A") turns out to contain the relatively active, gene rich regions of the chromosome while, the other ("B") is generally inactive and gene poor (5)[. We applied E-PCA to this data to study correlated structural fluctuations across the snapshots of chromosomes in different cells.

For Chr21, 34 TAD positions were measured for each cell (*N* = 34). We first constructed a matrix of pairwise distances between each combination of TADs along the chromosome, giving *m* = *N*(*N* − 1)/2 = 561 pairwise distance variables, since the *N* diagonal variables *u_ii_* are absent (always 0) in the distance matrix. We could further convert this into a contact matrix by defining a minimum distance that should be called a "contact", but here, we constructed the covariance matrix of distances across all 47 cells *C_ijkl_* = 〈( *l_ij_* – 〈*l_ij_*〉) (*l_kl_* – 〈*l_kl_*〉)〉, is the distance between TAD *i* and *j*, and 〈*l_ij_*〉 its ensemble average. We performed PCA on this covariance matrix of distances. For comparisons, we also performed I-PCA, MI-PCA, and Cartesian coordinate PCA (Supplemental Information) besides E-PCA.

How quickly the top eigenvalues of the covariance matrix decrease with the ranking index provides an overall idea of how degenerate the dataset is. If a conformational ensemble has only a few major modes of fluctuation, the first few eigenvalues will dominate the distribution and vice versa, if many eigenvalues are nearly equally high, we can conclude that there are many different motions in the ensemble of structures. The eigenvalues of this and other systems we examine in this study are shown in Fig. 2. To facilitate comparison across different types of matrices, we normalized these traces by the sum of the eigenvalues. Specifically, for this Chr21 microscopy data (3), we have examined the eigensystems of E-PCA, I-PCA, and Cartesian covariance matrices. Regardless of the method used, all eigenvalues drop to zero after the 46th eigenvector, reflecting that we have only 47 conformations (cells). An ensemble of *T* conformations generally can contain no more than *T* – *1* independent fluctuation modes around mean. Although we focus on reporting the results of Chr21 below, our study of chr22 TAD data revealed similar features (such as those shown in Figure 2) which demonstrated the robustness of our conclusions.

**Figure 2:**
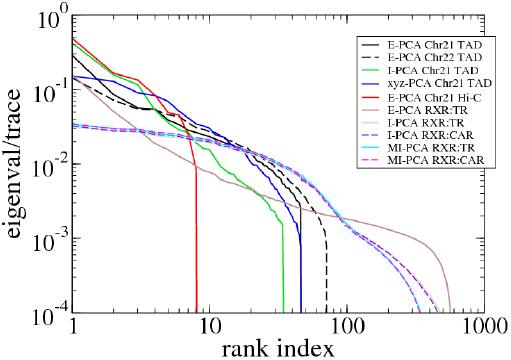
The eigenvalue distribution for various covariance matrices studied in this work. The main systems we consider (TAD imaging for chromosome 21, Hi-C from chromosome 10, and the protein complex RXR:TR) are shown in solid lines and comparison systems (TAD imaging from Chr22 and the protein complex RXR:CAR) are shown in dashes.

The top three principal components (PCs, eigenvectors of the covariance) of EPCA are displayed in a two-dimensional symmetric matrix known as displacement matrix *d_ij_*, as shown in Fig. 3(a-c). The displacement matrix has the same size (*N* × *N*) as the mean contact matrix, but they have different meanings. The elements of mean contact matrix <*u_ij_*> is never negative, while the displacement matrix *d_ij_* ~ *δu_ij_* = *u_ij_*−<*u_ij_*> can be either positive or negative. Each displacement matrix represents a specific mode of fluctuation around the mean. Each conformational ensemble only has one mean contact matrix but display many orthonormal modes of fluctuation. The displacement matrix shows how each explicitly expressed contact contributes to the given mode of fluctuation. When a particular contact, say one between *i* and *j*, shows strong blue (a highly positive value of *d_ij_*), the dynamics of that contact are highly important for this mode of fluctuation. Additionally, the contact dynamics of all blue regions are correlated, i.e., the contacts form and break in sync with each other. Similarly, the strongest red (negative) regions are also highly correlated with each other in their fluctuations, but opposite (anti-sync) from the blue regions. For example, if one red contact is formed, other red contacts are likely to be formed too, while blue contacts are likely to be broken. The corresponding three-dimensional rendition of eigenvectors is shown in Fig. 3(d-f) where strong fluctuation relationships between different parts of chromatin are rendered as colored cylinders. Note that we choose to use the 3D coordinates of the first cell to display TAD positions instead of an average (see Supplemental Information for further explanation). Both the matrix and 3D rendering of the E-PCA eigenvectors provide essential information on the biopolymer's structural changes as "concerted" modes of contact forming and breaking, or, in the current case using a distance matrix, positions moving toward and away from each other. In contrast, the eigenvectors of I-PCA analysis can be expressed as a bar graph (or a curve), such as MI-PC1 shown in Fig. 3a as a bar-graph label, Fig. 5(a-c), and Fig. S1 in SI, since each eigenvector of (M)I-PCA contains only N numbers. Particularly, in MI-PC1 of chromosomes, the most extreme positive and negative values can be used to indicate spatially segregated genomic bins, e.g., regions with positive values tend to associate with other positive regions.

**Figure 3:**
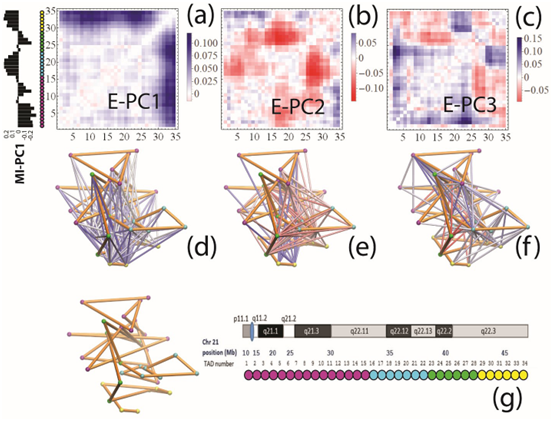
The E-PCA results for TAD imaging Chr21 system. The displacement matrix view of PC1, PC2, PC3 of E-PCA are shown in (a-c). PC1 from MI-PCA on the same system is shown as a bar-graph at left and TAD elements are colored according to general domains detected by E-PCA. The corresponding 3D representations are shown in (d-f), where the values of displacement matrix elements *b_ij_* are used to color cylinders that connect TADs i and *j.* Less significant relationships (cutoff is *|b_ij_|<* 0.06) are not shown. The reference 3D path of the 34 TADs (the configuration detected for a chosen cell, cell #1) is shown in (g) along with a reference for the genomic position of each TAD probe. Explanation was made in SI for the selection of cell #1 as the reference.

The dominant eigenvector PC1 of chr21, E-PC1, shows a prominent and simple inter-domain motion (association and dissociation) between two large-scale domains (TAD 1-28 and TAD 29-34), indicated by the strong blue region formed in Figure 3(a). The intradomain motions are primarily absent with small anticorrelated features (subtle red regions), i.e., when each domain is slightly more packed (indicated by the shortening of the distances between intradomain TADs, subtle red), the two domains dissociate (indicated by the lengthening of the distances between interdomain TADs, strong blue), and vice versa. The matrix for eigenvector PC2 in Fig. 3(b), on the other hand, shows intricate interactions between four independent regions (Domain 1: TAD 1-15, which belongs to the “B” compartment, colored by magenta; Domain 2: 16-22, “A”, cyan; Domain 3: 23-28, “B”, green; and Domain 4: 29-34, “A”, yellow) defined by their correlated dynamics. Interestingly, these domains uncovered in the correlated dynamics of interactions generally correspond well with the average domain compartmentalization identified by the top PC of MI-PCA (shown as a bar graph label in Fig. 3a). Indeed, the interactions contributing most strongly to PC2 (darkest red) are between neighboring domains, 2 and 3, that are, on average, spatially separated from each other in the “A” and “B” compartments respectively. When these two domains (2–3) move closer together, so does another pair of domains (1–2). The first two PCs suggest that the dynamics largely comes from the inter-compartment contacts (“A”-“B” type of contacts, such as 1-2, 2-3, and 1-4) with relatively little movement contribution from intra-compartment contacts (“A”-“A” or “B”-“B” type, such as 1-3).This might be thought of as a concerted “stretch and compress” motion within the first three regions of TADs. When the three domains stretch further away from each other, an anti-correlated interaction tends to be formed between the telomeric domain (TAD 29-34) and small local segments of the three domains (blue regions of the heatmap). Conversely, when the first three domains become aggregated, the last domain wants to break away.

In Fig.3(c), the mode of motion reported by eigenvector PC3 is much more localized. Similar to the interpretation of PC2, the classification of four regional domains (TAD 1-15, 16-22, 23-28, 29-34) can be utilized to describe these interactions. The third domain (TAD 23-28) is observed to form contacts with the first (TAD 1-15) and fourth domain (TAD 29-34), while the major parts of the second domain (TAD 16-22) migrate away from those two domains (first and fourth domains).

Overall, this E-PCA method reveals information about the dynamics of chromosome compartmentalization within individual cells, rather than just reporting an average spatial segregation. It allows us to begin to address persistent questions within the chromosome conformation field, such as how certain interactions or folding patterns relate to one another. These correlated movements could be related to the fact that different genes can be regulated in sync. To alleviate the concern that our results could be affected by the small sample size of chromosome conformations, we also checked the robustness of the E-PCA results by splitting 47 sampling points (individual cells) into two halves (first 23 and last 24) and investigating the level of the convergence, as discussed in SI. The results show the dot product of the normalized top eigenvectors is 0.834, which indicates the motions detected are not due to random noise.

### Key differences in 3D chromosome structure between cell types are captured by explicit contact correlation analysis

The above analysis shows that E-PCA holds promising potential for analyzing numerous chromosome contact maps (distance information) from individual cells. However, population-averaged Hi-C data is the only available structural information in many situations. Can contact PCA help us detect chromosome conformation changes using a set of average contact maps from different cell types or conditions? MI-PCA has been used toward this goal, and, by comparing vectors of PC1 values from MI-PCA, previous work has identified regions of chromosomes that, for example, switch their association from the “A” compartment to the “B” compartment on average in different cell types (9, 23). To compare changes in all interactions across Hi-C data from different cell types, rather than first reducing the Hi-C data to a representative single vector, we applied E-PCA to Chr21 and the first 17 Mb of Chr10 across Hi-C data from nine related human blood cell types (Figure 4a). We chose this region of Chr10 as a representative example from a mid-size chromosome with no major repetitive regions and strong A/B compartmentalization. We also performed analysis on TAD-averaged Hi-C data from Chr21 as a more direct comparison to the results from the imaging analysis. We focus on the results from Chr10 here, while the details of Chr21 analyses and the comparison with TAD imaging results are shown in SI. Specifically, the nine different cell types selected from the blood cell lineage were: 1. Neutrophil, 2. Monocyte, 3. M0 macrophage, 4. Naïve B-lymphocyte, 5. Naïve CD4+ T-lymphocyte, 6. Naïve CD8+ T-lymphocyte, 7. Lymphoblast cell line GM12878, 8. Erythroblast, 9. Megakaryocyte (21). Hi-C data was binned at 250 kb, to emphasize the compartment level of genome structure, and processed and normalized as previously described ((8, 24) and SI) The choice of chromosomal region and bin size here was chosen as a representative example of the application of this method. Other regions or levels of resolution could be chosen based on the particular biological focus or genes of interest.

**Figure 4:**
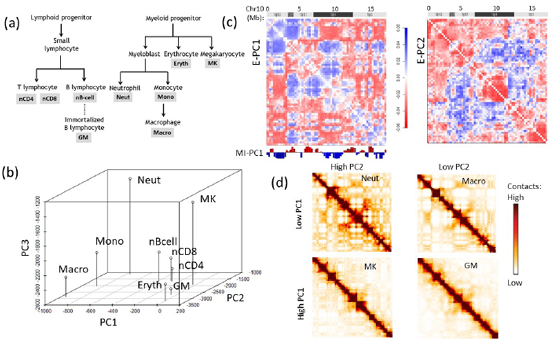
The E-PCA results for the Hi-C data from 9 different blood cell types. (a) The blood cell lineage from which the Hi-C data are derived, showing developmental relationships. Abbreviations used in subsequent panels are defined. (b) The projection of each cell type's Hi-C dataset onto E-PC1, 2, and 3. (c) The displacement matrix for E-PC1 and E-PC2, showing the 17 Mb region of chromosome 10 from which the sample Hi-C data was drawn. (d) Representative Hi-C contact matrices for cell types with high and low E-PC1 or PC2 projections, visibly showing the contact pattern differences between these categories of cell types.

In the current case, the eigenvectors show the dominant chromosome conformational differences across cell types, whereas in the first example, the conformational differences occur within an ensemble of cells of the same type. Since there are only 9 cell types, thus number of contact matrices, T = 9, only the top 8 eigenvectors and corresponding eigenvalues are nontrivial. Still, the statistics for this eigensystem is robust, since each Hi-C data point is not obtained from a single instant snapshot of the chromosome, but rather from an ensemble of millions of cells.

The results of the E-PCA analysis reflect the major features of the chromosome folding that distinguish one cell type from another. The projection of each cell type onto PC1, 2, and 3 in a 3D space nicely reflects known relationships and differences between cell types (Fig. 4b). From this projection alone, we can see that PC1 tends to segregate the myeloid lineage cells (macrophage, monocyte, and neutrophil; all have low PC1 projections) from the lymphoid lineage cells (which have higher PC1 projections). PC2 and PC3 further segregate within these major classes, distinguishing, for example, macrophages from neutrophils. Meanwhile, highly related cell types like nCD8 and nCD4 cluster near each other on all three PC axes. The displacement matrices for PC1 and PC2 (shown in Fig 4c, analogous to Fig 3a-c) show the interaction patterns that most distinguish these sets of cell types. Similar to the observations from the TAD imaging data above, the major features of PC1 relate to the A and B compartment segregation identified by MI-PCA. But, again, unlike MI-PCA, which focuses on the average associations of a genomic region with A or B and how that mean association changes between cell types, E-PCA reports on the *correlated* changes in the A/B compartmentalization of regions across the cell types. For E-PC1, the strongest positive values represent interactions between A and B compartment bins while the strongest negative values often represent interactions between regions of the same compartment identity. This result suggests that the strongest differentiator between cell types is the strength of compartment segregation. Indeed, representative contact matrices from cell types with high and low PC1 projections (Fig. 4d) show that cells like neutrophils and macrophages have a stronger segregation of A and B compartment regions (seen as a plaid pattern in the contact map) compared to cells like megakaryocytes and GM12878 lymphoblasts. E-PC2 shows that beyond strength of compartmentalization overall, there are more specific patterns of interaction within these broader domains that further distinguish between cell types. For example, across cell types, higher local interactions within 10p14 correspond to lower distant interactions between this region and neighboring regions, and vice versa.

### Implicit contact correlation analysis reveals consensus features of protein conformational ensembles

Besides using contact analysis for characterizing chromosome conformations, this type of analysis has been prominent for studying protein folding and structures. As noted above, E-PCA (four-point correlation analysis) was used previously for studying protein conformation dynamics and I-PCA (particularly, MI-PCA) for chromatin structural analysis. Here, we demonstrate what I-PCA can reveal about protein structure and/or dynamics. It is important to point out that I-PCA is beyond a two-point correlation analysis, since the correlation examined using I-PCA is not about the direct contacts formed between *i* and j. Although the size covariance matrix of I-PCA is, *N* × *N*, the computational complexity of I-PCA is not simply *0*(*N*^2^) but rather 0(*N*^3^), since each matrix contributes N sampling points (rows). I-PCA considers two types of deviations between samples: (i) a static one that emphasizes the difference between different rows of either of the same matrix or different matrices (snapshots), and (ii) a truly dynamic one, the variance of the same row in different matrices. The former term dominates over the latter, linearly increasing with the matrix size *N.* It is important to point out that MI-PCA only captures the static structure.

Two sets of molecular dynamics simulations of protein complexes were used in this study. The system setup details have been reported previously for E-PCA analysis (16, 17). Here, we focused on conformations from a long-time simulation of the wild-type complex(19). Both protein systems we considered contain a dimer of the ligand binding domains of nuclear hormone receptors. One system is a dimer complex between retinoid receptor (RXR) and thyroid receptor (TR), while the second system is a complex between same RXR and another nuclear receptor, constitutive androstane receptor (CAR). Including associated ligands, the RXR(9c):TR(t3) system has total N=493 residues with the internal indices as the following: RXR=1-232; TR=233-491; 9c=492; t3=493 (unless specified otherwise). Here ligands *9c* and *t3* are the corresponding ligands for RXR and TR respectively. Similarly, the complex RXR(9c):CAR(tcp) has N=476 with the following breakdown: RXR=1-232; CAR=233-474; 9c=475; tcp=476. Each simulation is 200 ns long with snapshots taken every 1 ps. Thus, T=200,000 conformations for each of the two complexes were converted into contact matrices for the I-PCA and E-PCA analysis.

Figure 5 shows the top four eigenvectors of I-PCA for the protein complex RXR:TR, both in the line representation and on a 3D cartoon representation of the protein complex, color-coded by the eigenvector value. The top eigenvector (Fig. 5a) shows a concerted anticorrelation between two monomers of the complex and naturally separate the complex as two halves (domains). One can also observe a certain level of symmetry between the monomers from PC1 of Figure 5. This is expected, given that the ligand binding domains of the nuclear hormone receptors share the same fold. The second eigenvector, PC2 also largely splits the complex into two halves, though the splitting planes are different. PC2 separates the bottom half (N-terminus, H1, H8, H9, half of H10, indicated by blue) and the top half (C-terminus, H6, H7, half of H10, indicated by red). Similarly, as shown in Fig. 5c, PC3 is yet another two-domain split while this time the front vs and the back. PC4 separates an interior core of the complex (center of H10 and H7) and the outside surface (the rest). Note that the top four eigenvectors are all large scale, global mode of fluctuation. To evaluate how robust our I-PCA results are, we also examined a second complex, RXR:CAR. One can see the corresponding top two eigenvectors show similarities overall, which indicates, at the large scale, the observed feature is robust. There are subtle differences observed: in PC2, for example, the absolute values of the eigenvector are larger for TR than RXR in the RXR:TR complex, while in complex RXR:CAR, CAR is more subdued than RXR. This would suggest that there is a stronger spatial separation of RXR and a more compact tructure for CAR.

**Figure 5:**
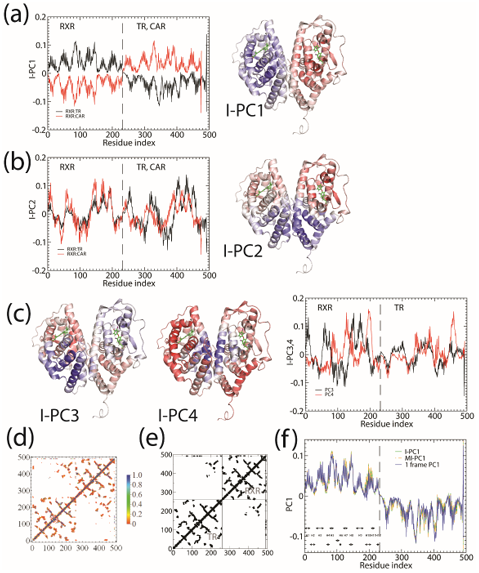
The I-PCA and MI-PCA results for protein conformations. (a) The top eigenvector, PC1 for RXR:TR and RXR:CAR are shown in the left panel, while the 3D view of the same information (for the RXR:TR case) is shown on the right panel. Here the values of the elements of eigenvector PC1 are displayed by color on a cartoon representation of the protein complex. (b) The corresponding information for eigenvector PC2. (c) The eigenvectors PC3 and PC4 for RXR:TR complex. (d) The top two eigenvectors PC1 and PC2 using MI-PCA.

MI-PCA only considers a single contact matrix and does not study the dynamic correlations between any contact events. As mentioned in the introduction, it has been the best-known analysis method to determine the spatial separation of chromosome compartments. It is interesting to see what MI-PCA can reveal about protein conformations. Here, we applied MI-PCA to protein systems using the mean contact map (shown in Fig.5d). For a comparison, we also tested the MI-PCA analysis using a single frame, replacing the mean contact matrix with one instant contact map (Fig. 5e). Surprisingly, the MI-PCA and, to a lesser extent, 1-frame PCA result in a very similar PC1 as was obtained using I-PCA on the full set of individual snapshots for the RXR:TR system, as shown in Fig. 5f. Similar results for other eigenvectors are shown in the Supplementary Information. Larger differences between I-PCA and the corresponding MI-PCA results were apparent for subsequent, high ranked eigenvectors; in particular, differences between MI- and I-PCA results were observed in the CAR part of PC2 for the RXR:CAR system, shown in SI. Similarity between I-PCA and MI-PCA, especially for the top eigenvectors, indicates the I-PCA analysis largely reveals the consensus features of the conformational ensemble, despite being a method analyzing covariance fluctuation and the inclusion of individual snapshots. This focus on consensus features arises from the fact that each row (or column) of the contact matrix is treated as an independent entity in this method. The total variation between rows of the same contact matrix is far more significant than the difference between the same row of different contact matrices–One can expect a high similarity between MI-PCA and I-PCA for a highly structured biopolymer, especially when *N* is large, but we expect that the level of similarity would decrease for semi-structured biopolymers such as chromosomes (from TAD data) and intrinsically disordered proteins.

Traditionally, E-PCA and Cartesian PCA include a PC projection analysis, i.e. render the original conformation sampling points using the newly found top PCs, e.g. the fc^th^ PC, the projection is *Σ_ij_d_ij_*^(*k*)^*U*_*ij*_(*t*). Each conformation of the biomolecule (snapshot) is displayed on the new coordinates spanned by the PCs, as previously demonstrated in Ref. (14–16, 25) and in the previous section of this current work for chromosomes. One can observe the fluctuation of individual conformations and the amplitude of fluctuation, where PCs themselves are normalized eigenvectors and do not provide overall amplitude. We tested PC projections for I-PCA (*Σ_i_d_i_*^(*k*)^*u_i_*(*t*)) and found that a direct application of PC projection for I-PCA leads to a largely "spreading-out" pattern, as shown in SI. One reason is that in I-PCA, each conformation (contact matrix) contributes *N* sampling points. For total *T* conformations, there are *N* × *T* points with two types of variances that showed up: “static” (variance between different rows) and “dynamic” (variance of the same row in different matrices) types. If one further averages the PC projections from these N points, we will obtain a largely trivial Gaussian distribution with very few features. Perhaps, a more meaningful way to display the variance is by focusing on the “static” component, i.e., focusing on PC projection of the MI-PCA, as shown in SI.

### Comparing E-PCA and I-PCA approaches and resulting properties of protein and chromosome structures

With the results we have obtained, we can make a more detailed comparison between the two methods being used. Based on how fast eigenvalues decay with increasing rank, we observe that I-PCA contains fewer modes of fluctuation whereas E-PCA typically has a slower decay and thus many fluctuation modes. This makes sense since E-PCA focuses on the explicit details of contacts being made. However, E-PCA has a faster initial drop at the top eigenvalues which means it contains fewer dominant modes, as shown in Figure 2. From a statistical analysis perspective, both methods study the same amount of information. E-PCA has more independent variables and less sampling points, while I-PCA has more sampling points (by dividing one contact matrix to *N* “pieces”), but has less stochastic variables and less explicit treatment of the contact information. The fact that I-PCA has previously been exclusively associated with chromatin structure analysis reflects the fact that initially, Hi-C data were scarce and little was known about the basic domains of the genome structure. Thus, it was useful in this system to first focus on the consensus features of an ensemble. However, with an increasing amount of data and higher resolution chromosome structures being characterized, E-PCA may be useful in many situations for chromatin systems. Conversely, as I-PCA mainly reveals the common features, it has not been applied to protein fields previously in which structures are often already known and the dynamic fluctuation is the focus. However, one can imagine I-PCA will have useful applications for semi-structured polymers where the ground state is less defined. For example, I-PCA is suitable for identifying protein domains and self-interacting regions from simulation data of intrinsically disordered proteins (26).

On the other hand, E-PCA explicitly tracks the correlated dynamics of polymer contacts, thus provides high resolution information and detailed correlated motions of the biopolymer and/or structure variation in the ensemble. It requires more data points than I-PCA, and thus a main drawback of E-PCA is the fast rise of the size of the covariance matrix *N*^4^. Such higher order correlation analysis requires more computational resources for the PCA data reduction task. For this reason, previous work has involved selecting dynamic contacts or coarse-graining contacts in the protein system to make the matrix size manageable. Thus, I-PCA, with reduced dimensionality, has its value in studying protein structures, especially for the large protein complexes which contain thousands of amino acid residues. I-PCA provides simple dichotomy of dynamic motions, separating two types of domains: local contact increasing (folding) and local contact decreasing (unfolding), but does not provide a detailed description of more complicated motions involving multiple domains as was seen in the example of E-PC2 of Chr21 using TAD data.

Regardless of the types of the covariance matrix being considered, it is always interesting to ask whether these eigenvectors represent true dynamic motions of the biopolymers under investigation whether these are simply a way of illustrating the difference between different conformations in the ensemble. The answer has nothing to do with a particular analysis procedure but rather depends on how the ensemble data has been generated. In the current work, the example using protein structure data is clearly an example of truly dynamic motion, because the underlying data are time-related snapshots of the protein conformation. In contrast, the second example involving different blood cells types is clearly not reporting "dynamics" but instead conformational differences between different states. The first example with TAD data is more ambiguous, as chromosome conformation differences between cells could reflect either heterogeneous stable conformations or conformations that interconvert within cells at a physiologically relevant timescale.

An attempt at directly comparing the dynamic motions of proteins and chromosomes is hampered by the inequivalent sampling data. Simply judging from the eigenvalue distribution, it would be tempting to conclude that the motion of chromosomes is largely “frozen” while proteins show a larger variety of modes of dynamics. However, two factors make this an unfair comparison: the amount of data (< 100 snapshots in chromatin systems versus > 100,000 in protein systems) used for the current work is highly discordant and the chromosome structure is likely much more hierarchical than the protein structure.

## Conclusions

As demonstrated from three distinct types of structural information (TAD imaging, Hi-C, and computer simulation) of biopolymers across scales, PCA of contact information can provide a powerful description of structural consensus and fluctuation of proteins and chromosomes. Different types of contact analyses appear to have a preferred scale: I-PCA is suitable to identify the overall consensus picture (ground state) such as domains, using large scale, low resolution data with fewer conformations in hand; whereas E-PCA highlights the major differences of an ensemble (the dominant fluctuations around the ground state) when sufficient data is available.

## ACKNOWLEDGMENTS

We acknowledge the computational support for protein simulation from allocations of advanced computing resources (STAMPEDE2 at TACC) provided by XSEDE. We thank Jacob Sanders and Rosela Golloshi for generating the GM12878 Hi-C data analyzed here. This study makes use of data generated by the PCHI-C Consortium. A full list of the investigators who contributed to the generation of the data is available in Ref.(21). Funding for PCHI-C project was provided by the National Institute for Health Research of England, UK Medical Research Council (MR/L007150/1) and UK Biotechnology and Biological Research Council (BB/J004480/1). The PCHI-C Consortium bears no responsibility for the further analysis or interpretation of these data, over and above that published by the Consortium.

